# Language gene polymorphism pattern survey provided important information for education context in human evolution

**DOI:** 10.1101/2022.10.31.514632

**Authors:** Wei Xia, Zhizhou Zhang

**Affiliations:** School of Languages and Literature, Harbin Institute of Technology, Weihai, China 264209; BIOX Biotechnology Center, Harbin Institute of Technology, Weihai, China 264209

**Keywords:** Language gene, single nucleic polymorphism, ancient genome, Nepal

## Abstract

Study on language gene polymorphism patterns (LGPP) across different populations could provide important information on human evolution. In this study, as a preliminary observation, we adopted 148 single nucleic polymorphism (SNP) sites from 13 language genes, each with 4-13 SNPs. These SNPs were screened across 112 whole genome sequences (including 59 ancient genomes ranging from 2000 BP to 120000 BP) from five continents (Africa, Asia, Europe, North America, and South America). We found that four distinct LGPPs featured across human evolution history, though it is still to decipher whether they correspond to the three batches of out-of-Africa ancient humans and modern human; Surprisingly, ten ancient samples from the small country, Nepal, contain all 4 basic LGPPs, suggesting that the southern foothills and nearby of the Qinghai-Tibet Plateau were likely an agglomeration place for ancient humans; Chinese samples also have 3-4 basic LGPPs. Of note, some types of Neanderthals and Denisovans possessed a LGPP almost the same as modern humans.

## Introduction

### The general landscape of human evolution has been clearly described, but many details keep obscure

Asia is an important continent for human evolution, and more ancient geologic evidences are always pursued to characterize complementary roles in this region. Specifically, Asia was connected with other continents, including Africa and America, in a long period of the old geologic time, during which some anthropoid would have a chance to move among several continents, especially from South Asia to Africa. By now, there are a plenty of ape species only found in Asia and Africa, and those more primitive apes are only found in South and Southeast Asia (not in Africa)(Besides, today’s habitat of anthropoid and chimpanzee shall be the places historical glacial climate was never able to destroy), strongly suggesting that apes in Africa came from Asia [1–4].Later on, apes in Asia and Africa evolved independently but with the same genome basis, making it certain that anthropoids in different regions always had the same taxonomic Family or Genus. Periodical glacial climate and other natural forces forced anthropoids to move between different regions and interact with each other, so as to render anthropoids similar evolution directions, until *Homo erectus* emerged in more than one places in the world. Apparently, *Homo erectus* in different regions had little reproductive isolation, and they spanned most of time with different evolutionary stages at the same time during the history of human being. However, it seemed that *Homo erectus* in Africa possessed the fastest speeds in brain and language evolution, and it was the *Homo erectus* in Africa pioneered several large batches of exodus, accelerating the broad blending together with many local extinctions of *Homo* species across different continents.

It is reasonable to decipher the origin of modern humankind through genetics, though other disciplines (for example, archaeology)also have their own unique advantages. Genetic technologies such as Y chromosomal haplogroups [5–6] and Human mitochondrial DNA haplogroups [7–9]can directly display evolution relationship among different populations, and found the main modern human populations all around the world originated from Africa. This result is not equal to that all humankind originated from the sole Africa; those elapsing human species always have their own places in the history of human evolution. If we perform genome sequencing for each person in the world, it is still possible to find someone who has no African origin, though such a person may only be found in a very remote corner in deep mountains and forests. Here is a logical loophole to tackle: the African anthropoids that evolved into *Homo erectus* may come from places other than Africa [1–4,10–11]. If the anthropoid ancestor of the third batch African *Homo sapiens* originated from South and Southeast Asia, the human evolution history will be rewritten.

Some Chinese scholars started to provide evidences pointing to a potential that China area might be the earliest birth place for humankind [10–11]. In fact, the time for the formation of East African Rift Valley was largely overlapped with that for the eminence of Qinghai-Tibet Plateau [12–13], while the latter also brought huge challenges to local apes in South Asia. It is postulated that both East African Rift Valley and Qinghai-Tibet Plateau can promote evolutionary opportunities for man-like apes to become *Homo erectus*. It is also postulated that anthropoids in South Asia and Africa had the same origin, and the origin place is likely just the South Asia/Southeast Asia [1–4]. Anthropoids in both places, under the stimulus of East African Rift Valley and Qinghai-Tibet Plateau, respectively, independently evolved into *Homo erectus* and gradually spread around the world. However, *Homo erectus* in Africa seemed to acquire quicker speed in evolution due to uncharacterized reasons, leading to the emergence of *Homo sapiens* first in Africa, not in other places. But the above postulations need future investigations to confirm.

### Language is a unique ability of humankind, and language evolution shall be a key step of human evolution

Language ability is a core feature of humankind. The evolution process of language ability witnessed continuous improvement of human brain intelligence and speech organs. This process took several million to tens of million years. The earliest record for apes is about 55 million years old [10–11]. A recent study indicated that *Sahelanthropus tchadensis* (a shah Ape at 7 million year BP) can walk upright [14]. Another study concluded that late-stage *Homo erectus* shall be able to use oral language fluently [15]. Modern studies confirmed that communication between apes mainly depends on sign language [16].Later on, it took millions of years for brain to evolve gradually in order to solidify sign gestures and body language into specific brain signals, until these signals were embodied as vocal sounds. Vocal sounds are produced by mechanical movements of specific muscles, and this explains why the language control region and the movement control region in the brain are highly overlapped [17].

After bipedalism, anthropoid cranial capacity [18-19,19a,19b] gradually became larger partly because of more cooked food and gene mutations. Larger brain had more capacity to compute complex signals, and eye-ear-mouth these organs were controlled more precisely to utter primitive languages. Movements of two hands also greatly boost the evolution process of human language, and as studied, communication between apes mainly depends on sign language [16]. Actually, language process was the same as quadruped locomotion in the context of organ movements in that language is just the result of eye-ear-mouth-etc organ movements. Language gene polymorphism shall witness the evolutionary process of language ability. During human evolution, eye-ear-mouth belonged to organs that had most mechanical movements, and were exactly most primitive motor organs. They got chances to connect with most neurons in the brain, thus paved a way for complex function development in both language and intelligence. These language and intelligence ability were essential to fabricate better stone-tools and to deliver tool-using skills accurately to offsprings. Besides, generations of anthropoid also had to spend a long evolution time to harmonize audio/video stimulation signals with oral muscle movements in the brain so as to possess a perfect language system (thus becoming *Homo sapiens*).

Language evolution spanned the process from anthropoid, anthropopithecus (man ape), pithecanthropus (ape man), *Homo erectus* to *Homo sapiens*. There should be several key breaking points for language evolution, but it is unknown whether the key points were located in anthropoid-ape-ape man-*H erectus* or in ape man-*H erectus*–*H sapiens* period [19,19a,19b]. Observation on language gene polymorphism patterns (LGPP) through the above species will help to answer the above question. In this study, due to the sample limitation, we only performed the LGPP screening among ancient human DNA and modern human DNA samples.

### The concepts of language gene and language gene polymorphism pattern

A gene functionally associated with language ability is called language gene. When a language gene is mutated, language ability will be compromised, slightly or heavily, in many different ways. By now there are 17 known language genes [15, 20–21], each with dozens to thousands of single nucleotide polymorphism (SNP) or single nucleotide variation (SNV). So 17 language genes would have hundreds to tens of thousands SNV to describe LGPP and its evolution. Such studies would alleviate some puzzles in human origin: Africa origin, multi-origin or East Asia origin? At least LGPP investigations will provide complementary results for human evolution conclusions derived from genetics studies.

## Methods

### Language genes and their SNPs

Language is an emergent complicated function of human being, though many other animals also have their own ‘languages’. If a gene mutation is statistically or experimentally associated with a certain language function loss, it would be called language gene. Language gene SNP data were all randomly selected for each gene in the dbSNP database: https://www.ncbi.nlm.nih.gov/snp/; Table 1 listed 13 language genes (as a preliminary observation, only 13 language genes were employed at the time the manuscript was written), and a total 148 SNPs from these 13 genes were selected for this study.

**Table 1.** Language genes employed in this study

|  | Name | Compromised ability when mutated(example) | References |
| --- | --- | --- | --- |
| 1 | FOXP1 | Expressive language | 22 |
| 2 | FOXP2 | Speech | 23 |
| 3 | CNTNAP2 | Early language development | 24-26 |
| 4 | TPK1 | Syntactic and lexical ability | 27-28 |
| 5 | DCDC2 | Reading, dyslexia | 29-31 |
| 6 | KIAA0319 | Reading, dyslexia | 24,29,32-33 |
| 7 | TM4SF20 | Language delay; communication disorder | 34 |
| 8 | FLNC | Reading, language | 35 |
| 9 | ATP2C2 | Memory | 36 |
| 10 | ROBO1 | Phonological buffer | 37-38 |
| 11 | ROBO2 | Expressive vocabulary | 39 |
| 12 | CMIP | Reading, memory | 24,29,36 |
| 13 | NFXL1 | Speech | 40 |

**Table 2.** Tested 148 SNPs of thirteen language genes

| Abbreviation | Language gene/SNP | Abbreviation | Language gene/SNP | Abbreviation | Language gene/SNP |
| --- | --- | --- | --- | --- | --- |
| ROBO-10 | ROBO1 rs34841026 | FXP1 | FOXP1 rs7638391 | CMI-1 | CMIP rs201316817 |
| ROBO-1 | ROBO2 rs11127602 | FOXP1-1 | FOXP1 rs76145927 | CMI-2 | CMIP rs183876152 |
| ROBO-2 | ROBO2 rs10865561 | FOXP1-2 | FOXP1 rs75214049 | CMI-3 | CMIP rs183075361 |
| ROBO-3 | ROBO2 rs5788280 | FOXP1-3 | FOXP1 rs17008544 | CMI-4 | CMIP rs114894868 |
| ROBO-4 | ROBO2 rs3923745 | FOXP1-4 | FOXP1 rs17008063 | CMI-5 | CMIP rs79979027 |
| ROBO-5 | ROBO2 rs3923744 | FOXP1-5 | FOXP1 rs11914627 | CMI-6 | CMIP rs74031247 |
| ROBO-6 | ROBO2 rs1163750 | FOXP1-6 | FOXP1 rs7639736 | CMI-7 | CMIP rs60152409 |
| ROBO-7 | ROBO2 rs1163749 | FOXP1-7 | FOXP1 rs1499893 | CMI-8 | CMIP rs57603843 |
| ROBO-8 | ROBO2 rs1163748 | FOXP1-8 | FOXP1 rs1053797 | CMI-9 | CMIP rs35429777 |
| ROBO-9 | ROBO2 rs1031377 | FOXP1-9 | FOXP1 rs144080925 | CMI-10 | CMIP rs34119643 |
| ROBO-11 | ROBO2 rs78817248 | FOXP1-10 | FOXP1 rs17008224 | CMI-11 | CMIP rs16955675 |
| ROBO-12 | ROBO2 rs144468527 | FOXP1-11 | FOXP1 rs147756430 | CMI-12 | CMIP rs2288011 |
| ROBO-13 | ROBO2 rs17525412 | FLN-1 | FLNC rs2291569 | CMI-13 | CMIP rs1187121850 |
| ROBO-14 | ROBO1 rs77350918 | FLN-2 | FLNC rs2291568 | ATP-1 | ATP2C2 rs78371901 |
| ROBO-15 | ROBO1 rs6795556 | FLN-3 | FLNC rs2291566 | ATP-2 | ATP2C2 rs74038217 |
| ROBO-16 | ROBO1 rs35456279 | FLN-4 | FLNC rs2291565 | ATP-3 | ATP2C2 rs62640935 |
| TM1 | TM4SF20 rs6724955 | FLN-5 | FLNC rs2291563 | ATP-4 | ATP2C2 rs62640932 |
| TM2 | TM4SF20 rs44675173 | FLN-6 | FLNC rs2291562 | ATP-5 | ATP2C2 rs62640931 |
| TM3 | TM4SF20 rs4675172 | FLN-7 | FLNC rs2291561 | ATP-6 | ATP2C2 rs62050917 |
| TM4 | TM4SF20 rs4673192 | FLN-8 | FLNC rs2291560 | ATP-7 | ATP2C2 rs16973859 |
| TM5 | TM4SF20 rs4438464 | FLN-9 | FLNC rs2291558 | ATP-8 | ATP2C2 rs13334642 |
| TM6 | TM4SF20 rs4428010 | FLN-10 | FLNC rs2249128 | ATP-9 | ATP2C2 rs4782970 |
| TM7 | TM4SF20 rs4408717 | FLN-11 | FLNC rs117864464 | ATP-10 | ATP2C2 rs4782948 |
| TM8 | TM4SF20 rs13415654 | FLN-12 | FLNC rs35281128 | ATP-11 | ATP2C2 rs2435172 |
| TM9 | TM4SF20 rs80305648 | FLN-13 | FLNC rs371111092 | ATP-12 | ATP2C2 rs247885 |
| TM10 | TM4SF20 rs137891000 | DCD-1 | DCDC2 rs35029429 | ATP-13 | ATP2C2 rs247818 |
| TPK-1 | TPK1 rs113536847 | DCD-2 | DCDC2 rs2274305 | KIA-1 | KIAA0319 rs138160539 |
| TPK-2 | TPK1 rs79464600 | DCD-3 | DCDC2 rs34584835 | KIA-2 | KIAA0319 rs117692893 |
| TPK-3 | TPK1 rs77358162 | DCD-4 | DCDC2 rs33943110 | KIA-3 | KIAA0319 rs114195393 |
| TPK-4 | TPK1 rs28380423 | DCD-5 | DCDC2 rs33914824 | KIA-4 | KIAA0319 rs699461 |
| TPK-5 | TPK1 rs17170295 | DCD-6 | DCDC2 rs9467075 | KIA-5 | KIAA0319 rs699462 |
| TPK-6 | TPK1 rs12333969 | DCD-7 | DCDC2 rs9460973 | KIA-6 | KIAA0319 rs699463 |
| TPK-7 | TPK1 rs6953807 | DCD-8 | DCDC2 rs3846827 | KIA-7 | KIAA0319 rs730860 |
| TPK-8 | TPK1 rs17170295 | DCD-9 | DCDC2 rs3789219 | KIA-8 | KIAA0319 rs10946705 |
| TPK-9 | TPK1 rs67644764 | DCD-11 | DCDC2 rs33943110 | KIA-9 | KIAA0319 rs75674723 |
| TPK10 | TPK1 rs77358162 | DCD-12 | DCDC2 rs190254728 | KIA-10 | KIAA0319 rs75720688 |
| NFX-1 | NFXL1 rs1964425 | CNTN-1 | CNTNAP2 rs1637842 | KIA-11 | KIAA0319 rs150584710 |
| NFX-2 | NFXL1 rs1822030 | CNTN-2 | CNTNAP2 rs1637841 | KIA-12 | KIAA0319 rs115399701 |
| NFX-3 | NFXL1 rs1822029 | CNTN-3 | CNTNAP2 rs1479837 | KIA-13 | KIAA0319 rs7770041 |
| NFX-4 | NFXL1 rs1812964 | CNTN-4 | CNTNAP2 rs1468370 | FOXP2-1 | FOXP2 rs10227893 |
| NFX-5 | NFXL1 rs1545200 | CNTN-5 | CNTNAP2 rs1062072 | FOXP2-2 | FOXP2 rs10244649 |
| NFX-6 | NFXL1 rs1440228 | CNTN-6 | CNTNAP2 rs1062071 | FOXP2-3 | FOXP2 rs12705977 |
| NFX-7 | NFXL1 rs1371730 | CNTN-7 | CNTNAP2 rs987456 | FOXP2-4 | FOXP2 rs61732741 |
| NFX-8 | NFXL1 rs1036681 | CNTN-8 | CNTNAP2 rs700309 | FOXP2-5 | FOXP2 rs61758964 |
| NFX-9 | NFXL1 rs978094 | CNTN-9 | CNTNAP2 rs700308 | FOXP2-6 | FOXP2 rs62640396 |
| NFX-10 | NFXL1 rs920462 | CNTN-10 | CNTNAP2 rs3194 | FOXP2-7 | FOXP2 rs73210755 |
| NFX-11 | NFXL1 rs147017712 | CNTN-11 | CNTNAP2 rs535454043 | FOXP2-8 | FOXP2 rs1058335 |
| NFX-12 | NFXL1 rs13152765 | CNTN-12 | CNTNAP2 rs2373284 | FOXP2-9 | FOXP2 rs61753357 |
| NFX-13 | NFXL1 rs34323060 | CNTN-13 | CNTNAP2 rs61732853 | FOXP2-10 | FOXP2 rs144807019 |
|  |  |  |  | FOXP2-11 | FOXP2 rs182138317 |

### Sample genome sequences

All genome sequences (Table 3) were downloaded from ENA database (https://www.ebi.ac.uk/ena/browser/). Total 112 whole genomes (including 59 ancient genomes) from 5 continents (Africa, Asia, Europe, North America, and South America) were collected, among which, there are 27 from EastAsia (China), 10 from Nepal, 12 from other SouthAsia countries, 20 from Africa, 28 from Europe, 9 from SouthAm, 2 from NorthAm, 2 from SEAsia, 1 from MAsia and 1 from WAsia.

**Table 3.**
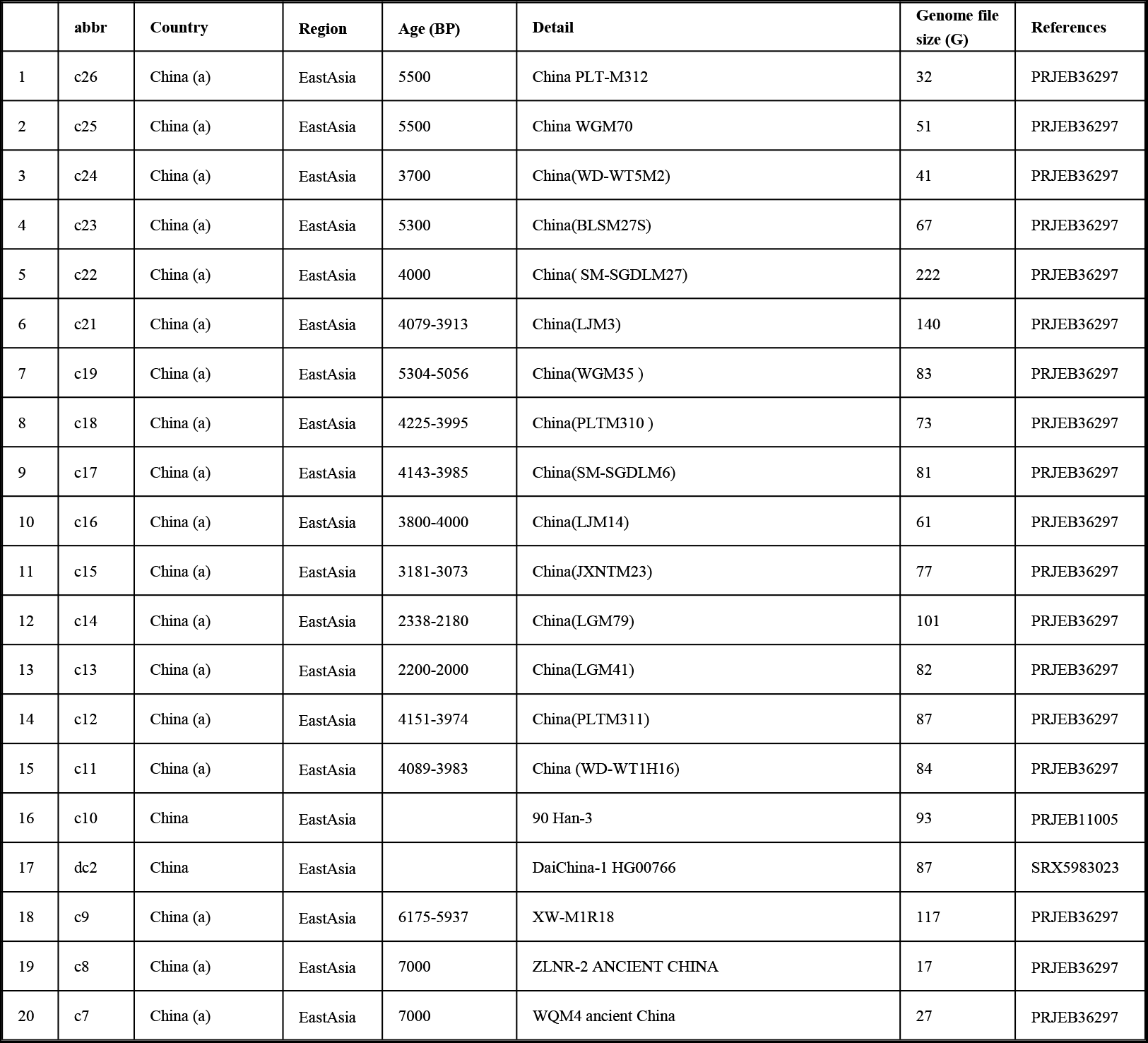
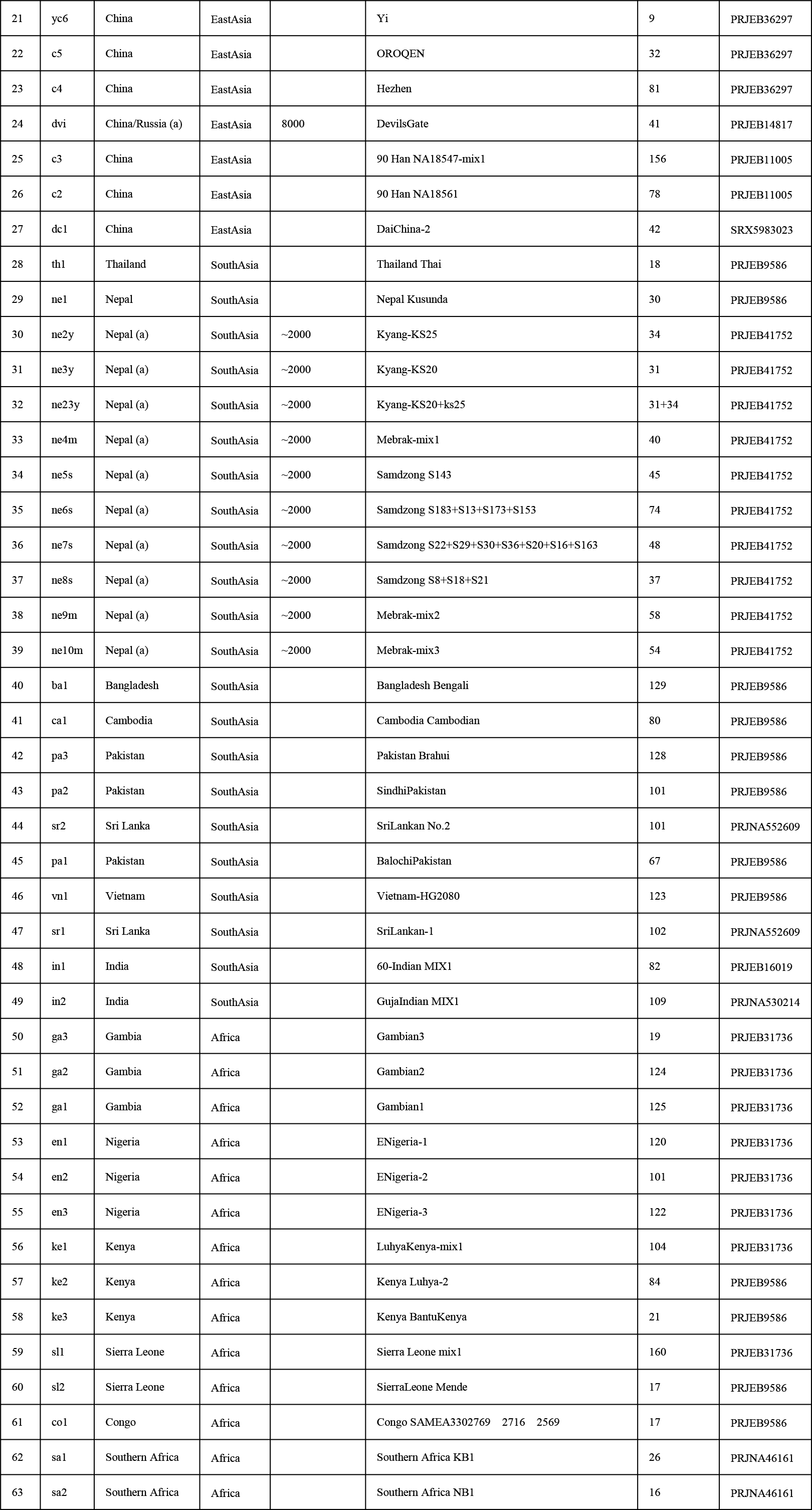
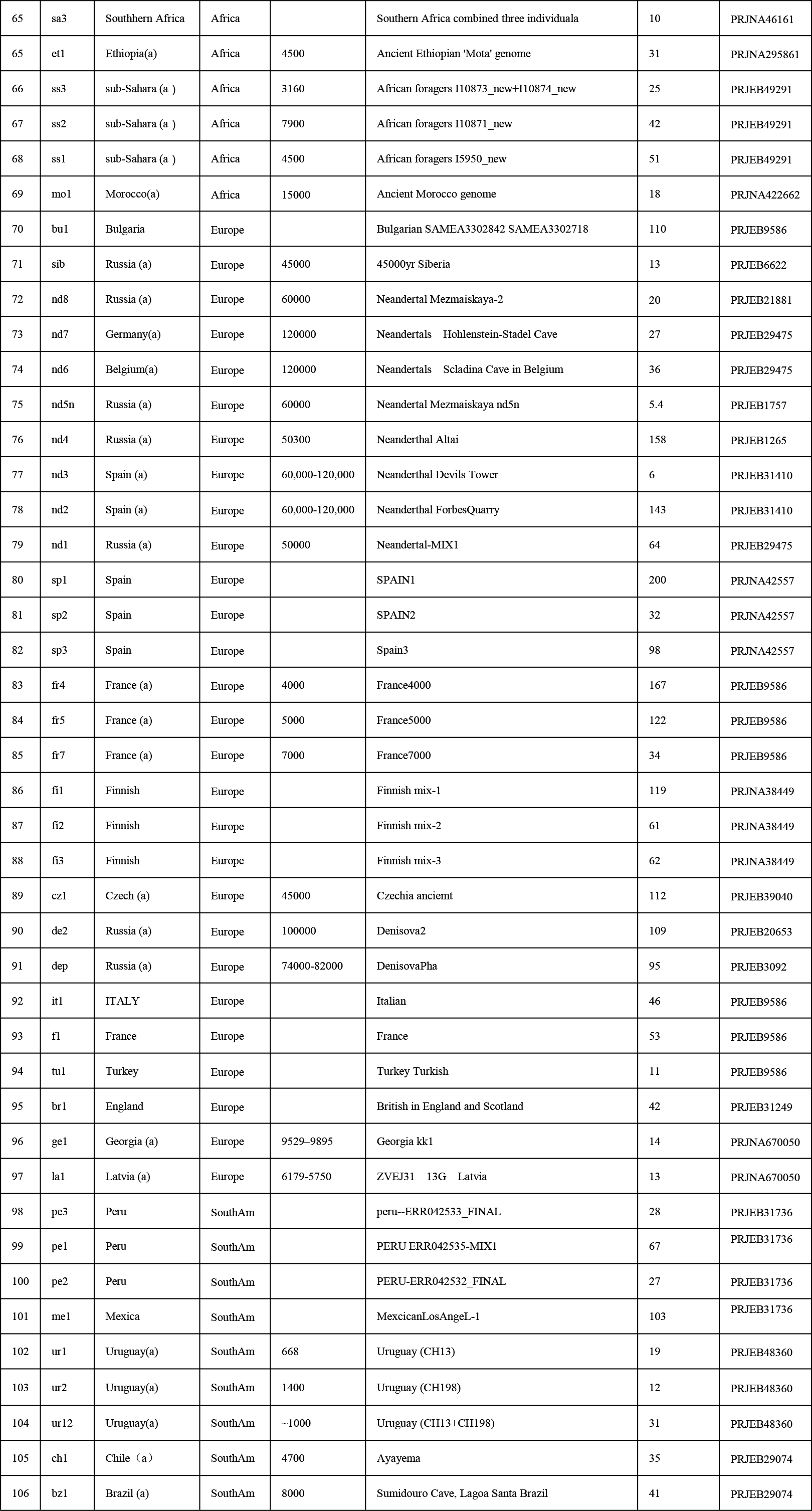
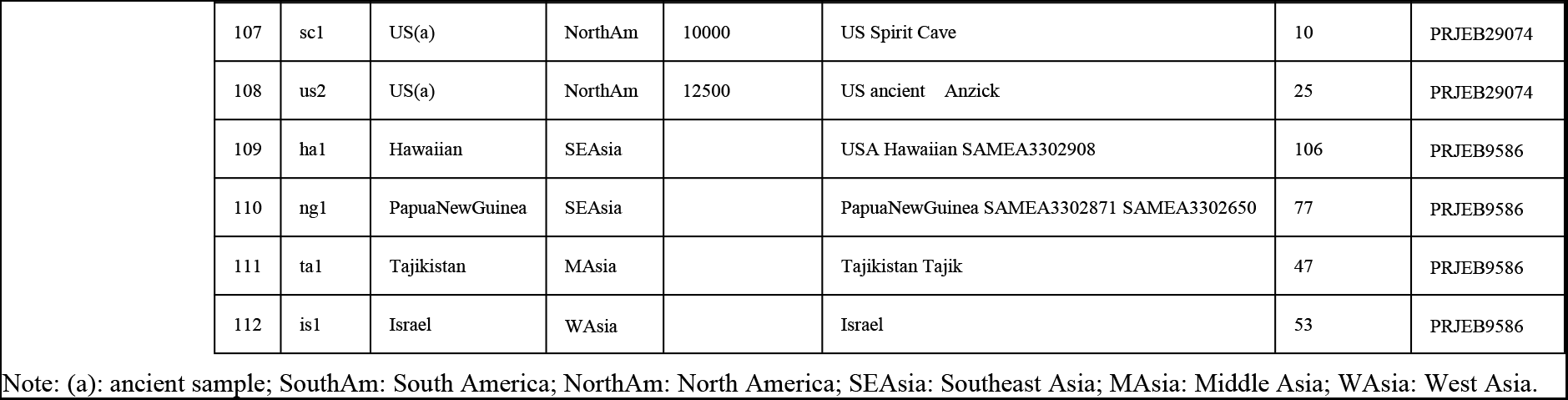
The 112 whole genomes employed in this study

### Sample SNP information abstraction and PCA analysis

The authors used 010Edit software to extract all 148 SNP information from each genome (supplementary file 1). In all 112 genomes, the sizes mainly range from 10G to 200G. Genomes less than 10G were neglected or only used as a reference. Principal Component Analysis (PCA) was performed using R packages FactoMineR, factoextra and ggplot2.The main R codes are listed as supplementary file 2.

## Result and Discussion

**Figure 1.**
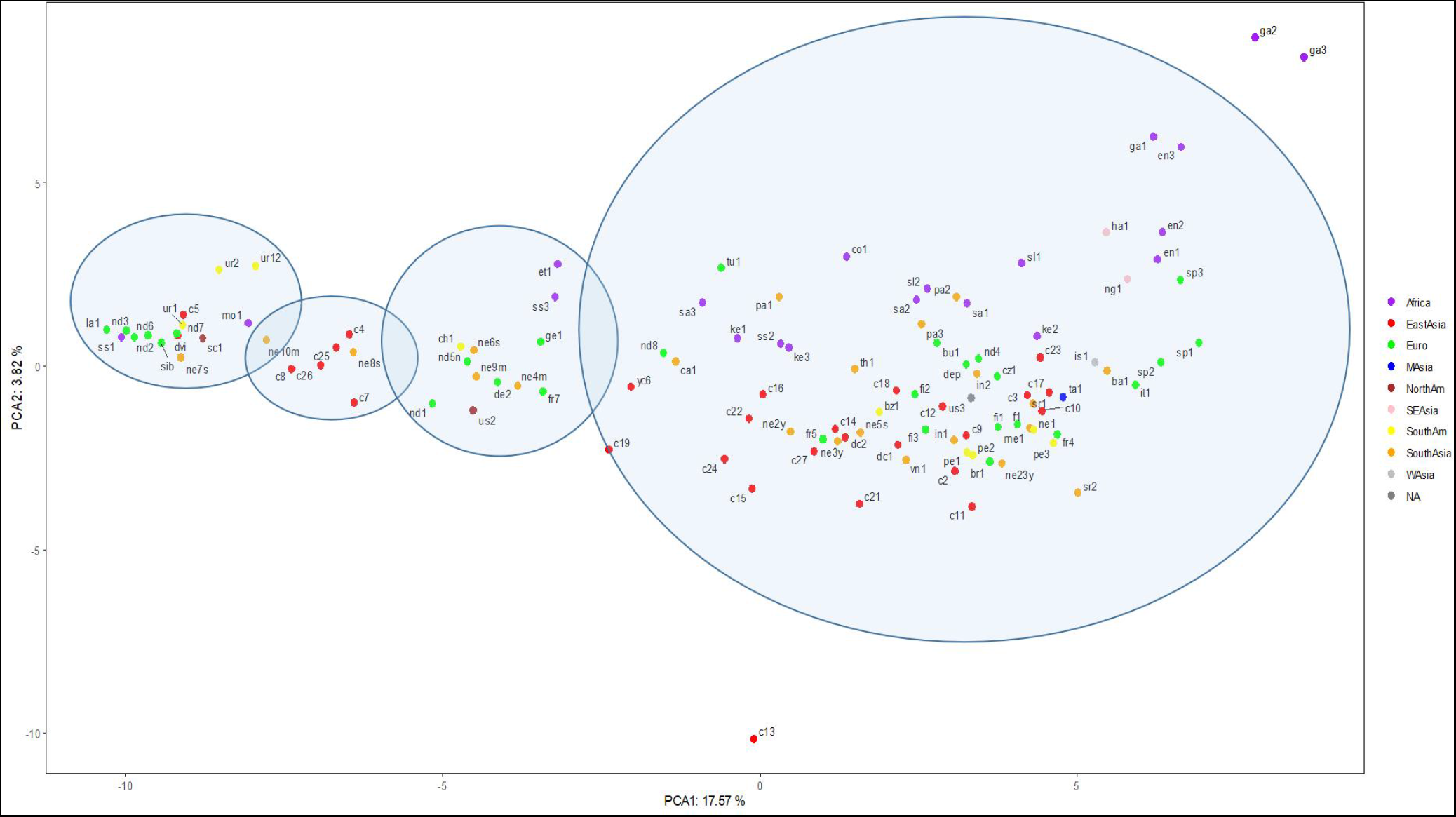
PCA results for 110 human genomes in the context of 148 language gene SNPs. From left to right, the four consecutive circles represent LGPP-1, LGPP-2, LGPP-3 and LGPP-4, respectively.

### Present data demonstrated Neanderthals possess three different LGPPs

Neanderthals’ LGPPs spanned three types except for LGPP-2. LGPP-2 was only represented by Nepalese and Chinese, and it was absent in Neanderthals in the samples used in this study. LGPP-2 may be only representing a batch of homonins directly that never touched the European land.

### LGPP-2 was absent in the American samples

Six American samples contain three different LGPPs: LGPP-1, LGPP-3 and LGPP-4. This suggests that at least two different batches (represented by LGPP-1 and LGPP-3)of human populations moved to the American land, consistent with present study results [41].If LGPP-1 and LGPP-3 represented those populations who successfully migrate into America, the Europe route, especially the whole Russia area, not likely the China route, was the one that mediated the migration. But interestingly, the results in the study [41] indicated that Chinese, Mongolian and Japanese were most likely similar to those ancient people in America.

### African samples possess unique and diverse LGPPs

LGPPs in African samples are very uncharacteristic in that most modern African didn’t well match the four LGPPs. One ancient African (ss1) sample matches LGPP-1, and mo1 lies between LGPP-1 and LGPP-2. Another two ancient African samples (et1 and ss3) are a little bit similar to, but apparently different from LGPP-3. It is also surprising that LGPP-1, LGPP-2 and LGPP-3 were not found in all modern African samples. This suggests that human evolution rate in Africa was likely more rapid than other continents, or the general diminishing rate of ancient populations was also faster in Africa. Some modern Chinese still possess very old LGPP, such as LGPP-1, indicating that some place in China holds a very stable environmental condition for a very long historical time, leading to an stable language gene polymorphism pattern(s) with little mutations.

### LGPP-3 was absent in Chinese samples

LGPP-3 corresponds to a group of human population containing nd1 (Neanderthals), nd5n (Neanderthals) and de2 (Denisovans), and it shall be this group that moved to France (fr7), Georgia (ge1), North America ((us2) and South America (ch1). This group was also similar to two ancient African people et1 and ss3. This group of people’s LGPP-3 was not seen in Chinese samples, but was observed in three Nepal samples (ne4m, ne6s and ne9m). This suggests that this group people shall come from Africa then to Europe. It also directly moved to Nepal from Africa, or moved to Nepal from Europe; but this group was not able to reach China.

### Nepal may be a crossroad where batches of ancient human populations stayed and left

Nepal is located at the southern foothills of the Qinghai-Tibet Plateau, and this place is relatively an enclosed area. It is not difficult to diffuse northwest or east from Nepal, but a little bit difficult to spread northeast where are China mountains. It will be most difficult to diffuse directly north through the Himalayan region as a directional barrier to gene flow [42]. The uplifting history of the Qinghai-Tibet Plateau demonstrated that the plateau was only about 1500 meter high 40 million years ago, probably providing conditions for mutual flow of apes between Africa and Asia. But it is extremely difficult to get LGPP information of ancient apes; what we can do right now is to analyze samples at hand. Chinese samples had three LGPPs while Nepalese samples had four, probably suggesting Nepal was easier to access than China for many ancient human species.

## Conclusion

Though the sample size is not large in this study, we still can get a glimpse of some important conclusions both for human evolution and for human evolution related education context. This study indicated that there are mainly four types of language gene polymorphism patterns (LGPP), though some populations especially modern Africans have apparent deviations from these four LGPPs (LGPP-1, LGPP-2, LGPP-3 and LGPP-4).The first three LGPPs were mainly related with ancient human samples, while the LGPP-4 is mainly related with modern human samples. Further investigations are needed to judge whether the first three LGPPs reflected the well-known three batches of out-of-Africa of ancient humankind [43–44]. Neanderthals possessed three LGPPs (LGPP-1, LGPP-3 and LGPP-4), while Denisovans only showed two LGPPs (LGPP-3 and LGPP-4; Only two Denisovan samples were used in this study). Interestingly, one Neanderthal sample (nd4)and one Denisovan sample (dep) had LGPP-4 like modern human, strongly suggesting that some Neanderthals and Denisovans are capable of speaking like modern human [15]. Surprisingly, ten ancient samples from the small country, Nepal, contain all 4 basic LGPPs, suggesting that the southern foothills and nearby of the Qinghai-Tibet Plateau were likely an agglomeration place for ancient humans. All the above preliminary observations have to be confirmed and elucidated with more language genes [35,45–48] and more SNPs (SNVs) in more samples in the near future.

## Supporting information

supplementary file 1--The 148 SNPs for 112 genomes

supplementary file 2-- R codes

## Acknowledgments

This study was supported by State Language Commission Research Grant (YB135-117) and National Research Center for Foreign Language Education Grant (ZGWYJYJJ10A042).

---

Supplementary file 1: The 148 SNPs for 112 genomes

Supplementary file 2: R codes

## References

[1] Erin Wayman. Out of Asia: How monkey and ape ancestors colonized Africa. SmithSonian, June 4, 2012

[2] Charles Q. Choi. Out of Asia: new origin proposed for humans, monkeys, apes. Live Science, October 27, 2010

[3] Jean-Jacques Jaeger, et al. Late middle Eocene epoch of Libya yields earliest known radiation of African anthropoids. Nature. 2010 Oct 28;467(7319):1095–8.

[4] Jean-Jacques Jaeger et al. New Eocene primate from Myanmar shares dental characters with African Eocene crown anthropoids. Nat Commun. 2019 Aug 6;10(1):3531.

[5] Mark A Jobling, Chris Tyler-Smith. The human Y chromosome: an evolutionary marker comes of age. Nat Rev Genet. 2003 Aug;4(8):598–612.

[6] P A Underhill, et al. Y chromosome sequence variation and the history of human populations. Nat Genet. 2000 Nov;26(3):358–61.

[7] Moreno, E. A war-prone tribe migrated out of Africa to populate the world. Nat Prec (2010). https://doi.org/10.1038/npre.2010.4303.1

[8] L Luca Cavalli-Sforza, Marcus W Feldman. The application of molecular genetic approaches to the study of human evolution. Nat Genet. 2003 Mar;33 Suppl:266–75.

[9] EKF Chan, et al. Human origins in a southern African palaeo-wetland and first migrations. Nature. 2019 Nov; 575(7781):185–189.

[10] Ni X, et al. The oldest known primate skeleton and early haplorhine evolution. Nature. 2013 Jun 6; 498(7452):60–4.

[11] Xijun Ni, et al. A euprimate skull from the early Eocene of China. Nature. 2004 Jan 1;427(6969):65–8.

[12] Xiong et al., The rise and demise of the Paleogene Central Tibetan Valley. Sci. Adv. 8, eabj0944 (2022)

[13] Duncan Macgregor. History of the development of the East African Rift System: A series of interpreted maps through time. Journal of African Earth Sciences. Volume 101, January 2015, Pages 232–252.

[14] Daver G, et al. Postcranial evidence of late Miocene hominin bipedalism in Chad. Nature. 2022 Sep; 609(7925):94–100.

[15] DG Hillert. On the Evolving Biology of Language. Front. Psychol. 2015 6:1796.

[16] Amy S Pollick, Frans B M de Waal. Ape gestures and language evolution. Proc Natl Acad Sci U S A. 2007 May 8; 104(19):8184–9.

[17] Dietrich Stout, Thierry Chaminade. Stone tools, language and the brain in human evolution. Philos Trans R Soc Lond B Biol Sci. 2012 Jan 12; 367(1585):75–87.

[18] P. V. Tobias. Cranial capacity in anthropoid apes, Australopithecus and Homo habilis, with comments on skewed samples. South African Journal of Science. 1968, 81–91.

[19] G. Lynch, S. Hechtel, D. Jacobs. Neonate size and evolution of brain size in the Anthropoid primates. Journal of Human Evolution (1983) 12, 519–522.

[19a] Marcia S Ponce de León et al. The primitive brain of early Homo. Science. 2021 Apr 9; 372(6538): 165–171.

[19b] Zachary Cofran. Brain size growth in Australopithecus. J Hum Evol. 2019 May;130:72–82.

[20] Jie Huang, Wei Xia, Hongrui Ji, Zhizhou Zhang. General correlation profile between the basic language parameters and language gene polymorphisms plus multiple edu-geo-cul-soc parameters of twenty-six countries. ICMET2022: Proceedings of the 4th International Conference on Modern Educational Technology May 2022 Pages 127–134.

[21] Zongxuan Liu, Wei Xia, Bo Sun, Changlu Guo, Zhizhou Zhang. Correlation analysis between language gene polymorphism and geography/society parameter from twenty-six countries. 2021, Research Square, DOI: https://doi.org/10.21203/rs.3.rs-960107/v1

[22] Bacon C & GA Rappold. The distinct and overlapping phenotypic spectra of FOXP1 and FOXP2 in cognitive disorders. Human Genetics, 2012,131(11):1687–9168.

[23] CS Lai et al. A fork-head domain gene is mutated in a severe speech and language disorder. Nature, 2001, 413(6855):519–523.

[24] Newbury, D. F. et al. Investigation of dyslexia and SLI risk variants in reading-and language-impaired subjects. Behav. Genet. 41, 90–104; (2011).

[25] Vernes, S. C. et al. A functional genetic link between distinct developmental language disorders. N. Engl. J. Med. 359, 2337–2345;(2008).

[26] Whitehouse, A. J., Bishop, D. V., Ang, Q. W., Pennell, C. E. & Fisher, S. E. CNTNAP2 variants affect early language development in the general population. Genes Brain Behav. 10, 451–456;(2011).

[27] Villanueva P et al. Genome-wide analysis of genetic susceptibility to language impairment in an isolated Chilean population. European Journal of Human Genetics, 2011, 19(6):687–695.

[28] Fattal I et al. The crucial role of thiamine in the development of syntax and lexical retrieval:a study of infantile thiamine deficiency. Brain, 2011, 134(6):1720–1739.

[29] Scerri, T. S. et al. DCDC2, KIAA0319 and CMIP are associated with reading-related traits. Biol. Psychiatry 70, 237–245;(2011).

[30] Deffenbacher, K. E. et al. Refinement of the 6p21.3 quantitative trait locus influencing dyslexia: linkage and association analyses. Hum. Genet. 115, 128–138; (2004).

[31] Schumacher, J. et al. Strong genetic evidence of DCDC2 as a susceptibility gene for dyslexia. Am. J. Hum. Genet. 78, 52–62;(2006).

[32] Paracchini, S. et al. The chromosome 6p22 haplotype associated with dyslexia reduces the expression of KIAA0319, a novel gene involved in neuronal migration. Hum. Mol. Genet. 15, 1659–1666;(2006).

[33] Francks, C. et al. A 77-kilo base region of chromosome 6p22.2 is associated with dyslexia in families from the United Kingdom and from the United States. Am. J. Hum. Genet. 75, 1046–1058;(2004).

[34] Wiszniewski W et al. TM4SF20 ancestral deletion and susceptibility to a pediatric disorder of early language delay and cerebral white matter hyperintensities. American Journal of Human Genetics, 2013, 93(2):197–210.

[35] Gialluisi A et al. Genome-wide screening for DNA variants associated with reading and language traits. Genes Brain and Behavior, 2014, 13(7):686–701.

[36] Newbury, D. F. et al. CMIP andATP2C2 modulate phonological short-term memory in language impairment. Am. J. Hum. Genet. 85, 264–272; (2009).

[37] Hannula-Jouppi, K. et al. The axon guidance receptor gene ROBO1 is a candidate gene for developmental dyslexia. PLoS Genet. 1,e50; (2005).

[38] Bates, T. C. et al. Genetic variance in a component of the language acquisition device: ROBO1 polymorphisms associated with phonological buffer deficits. Behav. Genet. 41, 50–57;(2011).

[39] StPourcain, B. et al. Common variation near ROBO2 is associated with expressive vocabulary in infancy. Nat. Commun. 5, 4831; (2014).

[40] Villanueva, P. et al. Exome sequencing in an admixed isolated population indicates NFXL1 variants confer a risk for specific language impairment. PLoS Genet. 11,e1004925; (2015).

[41] N. von Cramon-Taubadel, A. Strauss, M. Hubbe, Evolutionary population history of early Paleoamerican cranial morphology. Sci. Adv. 3, e1602289 (2017)

[42] Tenzin Gayden, et al. The himalayas as a directional barrier to gene flow. Am. J. Hum. Genet. 2007; 80:884–894.

[43] López et al. Human dispersal out of Africa: a lasting debate. Evolutionary Bioinformatics 2015:11(s2) 57–68 doi: 10.4137/EBo.s33489.

[44] Chris Stringer Julia Galway-Witham. When did modern humans leave Africa? Science, 359 (6374)

[45] Pedro Tiago Martins, Maties Mari‱ and Cedric Boeckx. SRGAP2 and the gradual evolution of the modern human language faculty, Journal of Language Evolution, 2018, 67–78.

[46] Lei Xing et al. Expression of human-specific ARHGAP11B in mice leads to neocortex expansion and increased memory flexibility. EMBO J. 2021 Jul 1;40(13):e107093.

[47] Taipale, M. et al. A candidate gene for developmental dyslexia encodes a nuclear tetratricopeptide repeat domain protein dynamically regulated in brain. Proc. Natl. Acad. Sci. USA 100,11553–11558; (2003).

[48] Paracchini, S. et al. Analysis of dyslexia candidate genes in the Raine cohort representing the general Australian population. Genes Brain Behav. 10, 158–165; (2011).

