## supplementary file 2-- R codes for "Language gene polymorphism pattern survey provided important information for education context in human evolution": supplementary file 2-- R codes.pdf

The main R codes are listed as supplementary file 2.

```
> library(FactoMineR)
> library(factoextra)
> library(ggplot2)
> country <- read.delim('C:/RBook/20220516fastqSNPdata.txt', row.names = 1, sep = '\t')
> country <- t(country)
> country.pca <- PCA(country, ncp = 2, scale.unit = TRUE, graph = FALSE)
> plot(country.pca)
> pca_sample <- data.frame(country.pca$ind$coord[ ,1:2])
> head(pca_sample)
> pca_eig1 <- round(country.pca$eig[1,2], 2)
> pca_eig2 <- round(country.pca$eig[2,2],2 )
> pca_eig1
> pca_eig2
> group <- read.delim('C:/RBook/group3.txt', row.names = 1, sep = '\t', check.names = FALSE)
> group <- group[rownames(pca_sample), ]
> pca_sample <- cbind(pca_sample, group)
> pca_sample$samples <- rownames(pca_sample)
> head(pca_sample)
> library(ggrepel)
> ggplot(data = pca_sample, aes(x = Dim.1, y = Dim.2)) +geom_point (aes(color = group), size = 3) +  scale_color_manual (values = c ('purple', 'red', 'green','blue','brown', 'pink','yellow','orange','grey')) + theme(panel.grid = element_blank(), panel.background = element_rect (color = 'black', fill = 'transparent'), legend.key = element_rect (fill = 'transparent')) + labs(x = paste('PCA1:', pca_eig1, '%'), y = paste('PCA2:', pca_eig2, '%'), color = '') + geom_text_repel (aes (label = samples), size = 3, show.legend = FALSE, box.padding = unit(0.25, 'lines'))
```
